# The effect of chronic and acute stressors, and their interaction, on gonadal function: an experimental test during gonadal recrudescence

**DOI:** 10.1101/156034

**Authors:** Mikus Abolins-Abols, Rachel E. Hanauer, Kimberly A. Rosvall, Mark P. Peterson, Ellen D. Ketterson

## Abstract

Organisms are expected to invest less in reproduction in response to a stressor, but theory predicts that this effect should depend on the frequency of stressors in the environment. Here we investigated how an acute stressor affected gonadal function in a songbird, and how long-term differences in the stress environment influenced these acute stress responses. We exposed male Dark-eyed Juncos (*Junco hyemalis*) either to chronic or minimal (control) disturbance during gonadal recrudescence, after which we measured baseline testosterone, testosterone after an acute handling stressor, and the ability to elevate testosterone in response to hormonal stimulation. In a 2x2 design, we then euthanized males from the two chronic treatment groups either immediately or after an acute stressor to investigate the effect of these treatments on the gonadal transcriptome. We found that chronically disturbed birds had marginally lower testosterone. The acute stressor suppressed testosterone in control birds, but not in the chronic disturbance group. The ability to elevate testosterone did not differ between the chronic treatments. Surprisingly, chronic disturbance had a weak effect on the testicular transcriptome, and did not affect transcriptomic response to the acute stressor. The acute stressor, on the other hand, upregulated cellular stress response, and affected expression of genes associated with hormonal stress-response. Overall, we show that both chronic and acute stressors affect reproductive function, and that chronic stress changes how acute stressors affect testosterone physiology. Our findings also suggest that acute and chronic stressors affect testes differently, and that gonadal function is relatively robust to long-term stressors.

**Summary statement:** An acute stressor downregulated testosterone production, but this effect was absent in chronically disturbed birds. The acute stressor had a strong effect on the gonadal transcriptome, whereas chronic disturbance had a negligible effect.

## 1. Introduction

It is well-known that stressors can have a profound negative effect on reproductive physiology and behavior (Chand and Lovejoy, 2011; Rivier and Rivest, 1991; Selye, 1946). This effect has been demonstrated in relation to a variety of stressors, including food limitation (Lynn et al., 2015), thermal stress (Hansen, 2009), and psychological stress (McGrady, 1984; Nargund, 2015). The main adaptive hypothesis for this suppressive effect states that reproduction is inhibited because the physiological, energetic, and behavioral components of reproduction are costly and may directly interfere with resource and time allocation to stress response (Breuner et al., 2008). Because the stress response is an integral part of self-maintenance and survival, the interaction between stress and reproduction constitutes an important aspect of the life-history trade-off between current and future reproduction (Wingfield and Sapolsky, 2003).

Life history theory predicts that the effect of stress on reproduction should depend on the costs and benefits of both of these functions (Wingfield and Sapolsky, 2003). For example, if future reproductive success (residual reproductive value) is expected to be low, succeeding at the current reproductive effort is crucial to the fitness of the organism. It is expected that under these conditions reproduction should be more resistant to the suppressive effects of stressors compared to cases when future reproductive success is expected to be high. Consistent with this prediction, stress has been shown to be less suppressive of reproduction in semelparous organisms and organisms that have short seasonal breeding seasons (Wingfield and Sapolsky, 2003). At an intraspecific level, stressors have been shown to have a reduced effect on reproductive behaviors in populations that inhabit high-disturbance urban areas compared to rural habitats (Abolins-Abols et al., 2016).

While these findings are consistent with the theoretical predictions, we lack experimental studies that explicitly test these predictions and study which conditions lead to changes in the suppressive effect of stress on reproduction. Revealing the conditions and mechanisms that mediate effects of stress on reproduction is important for a better understanding of reproductive physiology and ecology of animals (Calisi et al. 2017), especially in the context of the rapid human-induced environmental change (Sih et al. 2011). Ultimately, such knowledge will aid in developing a predictive understanding of how and why individuals or populations respond differently to stressors.

We are addressing this question by focusing on the gonad, as one of the most important reproductive organs. In vertebrate males, testes produce sperm and androgen hormones, such as testosterone. Testosterone serves multiple important roles in reproduction: it regulates the development of the sexual phenotype (Hau, 2007), and is important in mediating behaviors necessary for reproduction, such as courtship (Fusani et al., 2014; Hutchinson, 1967) and territorial behavior (Soma, 2006; Wingfield et al., 1987). Testosterone synthesis is largely regulated by the hypothalamic-pituitary-gonadal (HPG) axis, where gonadotropin releasing hormone (GnRH) from hypothalamus causes the secretion of luteinizing hormone (LH) from the pituitary, which in turn upregulates synthesis of testosterone by the Leydig cells in gonads (Farner and Wingfield, 1980; London et al., 2006). However, testosterone synthesis by gonads has also been shown to change in response to local signaling factors independent of brain signaling (Nogueiras et al., 2004; Tena-Sempere et al., 2002). Testosterone levels can be suppressed by both psychological (Deviche et al., 2010; Moore and Mason, 2000) as well as physical stressors (Lynn et al., 2015; Nelson et al., 1989). This decrease may be mediated via suppressive effects of stressors on GnRH or LH secretion (Breen and Karsch, 2006; Breen et al., 2007; Nikolarakis et al., 1986) or by direct action of components of the physiological stress response, such as the hypothalamic-pituitary-adrenal (HPA) axis, on gonads (Lynn et al., 2015; McGuire et al., 2013).

Gonads have been shown to be especially sensitive to stressors and other perturbations during development (Guillette Jr et al., 1994; Rhind et al., 2001; Zambrano et al., 2014). For example, maternal stress during fetal gonadal development can result in reduced gonadal size (Dahlöf et al., 1978), and exposure to glucocorticoids during fetal development has been shown to reduce testosterone synthesis capacity *in vitro* (Page et al., 2001) and delay the onset of puberty (Smith and Waddell, 2000). Such developmental programming has important implications for pathology and reproductive rates in the wild (McMillen and Robinson, 2005; Sheriff et al., 2010). If gonadal responses to developmental stress are adaptive, then the effect of stressors on gonadal function should conform to the life history expectations, as outlined above, wherein gonads are expected to be less sensitive to stressors in animals undergoing development in a high-stress environment compared to animals developing in a benign environment.

In this study we experimentally investigated how acute and chronic stressors experienced during seasonal gonadal recrudescence (development) change gonadal function in Dark-eyed Junco (*Junco hyemalis*), a small passerine. In particular, we asked if chronic disturbance and acute stressors, and their interaction, changed baseline testosterone levels, the overall ability to elevate testosterone following stimulation with exogenous GnRH, testicular sensitivity to regulation by HPG and HPA axes, and testicular transcriptome in general. In general, we expected both types of stressors to reduce reproductive function in animals with actively recrudescing gonads, but that animals in high disturbance environment would show a reduced sensitivity of reproductive physiology to acute stressors, compared to animals in a low disturbance (control) environment. Specifically, we predicted that both chronic and acute stressors should decrease testosterone levels, but that the suppressive effect of an acute stressor on testosterone synthesis would be dampened in chronically disturbed animals. We further predicted that both chronic and acute stressors would negatively affect genes associated with testosterone production and spermatogenesis, up-regulate expression of genes involved in cellular and hormonal stress response, but that the effect of acute stressor on the gonadal transcriptome would be dampened in the chronically disturbed animals. More specifically, we expected that stressors would down-regulate steroidogenesis gene expression and up-regulate receptors for stress hormones implicated in regulation of gonadal physiology, such as glucocorticoid-receptor (GR)(McGuire et al., 2013) and gonadotropin-inhibitory hormone receptor (GnIHR)(McGuire and Bentley, 2010).

## METHODS

### 2.1 Capture and housing of study organisms

We captured 36 wintering male Dark-eyed Juncos *Junco hyemalis hyemalis* in Bloomington, Indiana, in December 2013 using baited mistnets and walk-in traps. We determined the sex of birds using plumage color and confirmed it with genetic sex markers (Griffiths et al., 1998). Before the experiment, birds were housed in free-flying groups in indoor aviary rooms, reflecting their natural winter flocking lifestyle. On January 15, 2014 we started to gradually (3 times per week) increase day length from natural Indiana winter photoperiod (10:44 daylight hrs, **Figure 1**). On January 27 males were transferred from free flying flocks to individual 2x2 ft metal cages, 9 cages per room, with visual access to other birds, including females. Cages contained food, water bowls, and perches. Food was provided *ad libitum* three times a week, and cages were cleaned once a week. When the experiment began on February 3, animals were experiencing 13:23 hrs of daylight, and by the end of the experiment on February 26, birds were exposed to 16:03 hrs of daylight, reflecting a summer-like photoperiod. Males were singing during the last week of the experiment.

**Figure 1:**
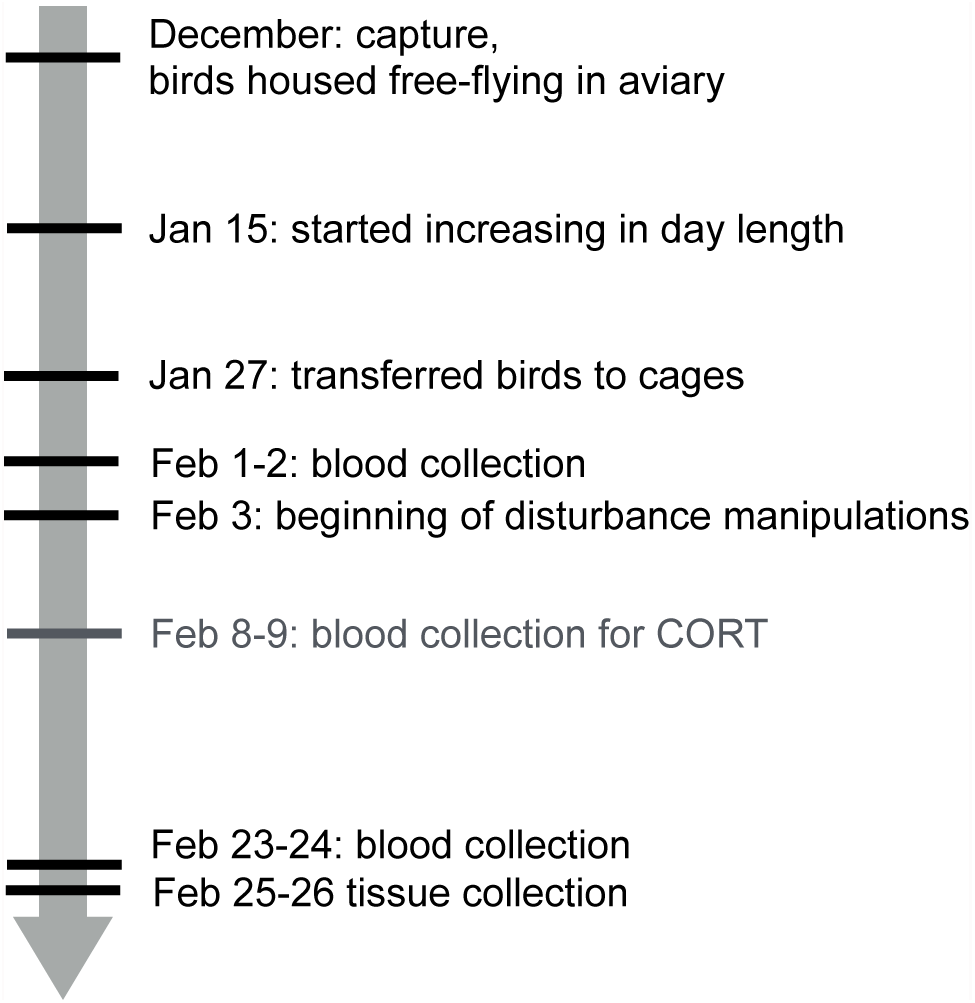
Timeline of the experiment

### 2.2. Chronic disturbance treatment

We randomly assigned half of the birds (n=18) to a chronic disturbance treatment, while the other half (n=18) were treated as controls. In the chronic disturbance treatment, birds were exposed to either physical disturbance (hand-waving, cage-tapping) or to a predator mount for 30 min four times each day for three weeks. During physical disturbances, observers (one per room) either waved their hand inside the cage, or tapped on the cage for 30 sec, after which the observer moved on to another cage in random order. For the predator stressor, we used taxidermic study skins of Cooper’s Hawk, Barn Owl, and Fox Squirrel, which were fixed to a tripod and left in the room by the observer. All these treatments had previously been validated in a pilot study and shown to result in an increase in corticosterone, a measure of a physiological stress response (Hanauer et al. in prep). In the pilot study, physical disturbance caused the highest increase in corticosterone, therefore birds in the chronic disturbance treatment received at least one of these disturbances each day (average: 2.5 physical disturbances each day). Otherwise, the choice of the stressors for each disturbance bout was random. The disturbance bouts occurred at least 60 min apart for each room and were administered during daylight hours. In the control group, we did not disturb the birds except for standard animal husbandry (see above).

### 2.3. Blood sampling and acute handling stressor

We took blood samples from each bird to analyze testosterone and corticosterone concentrations before (February 1-2) and after the three weeks (February 23-24) of chronic disturbance or control treatment. In addition, blood was taken from birds one week after the beginning of treatment for analysis of corticosterone levels (February 8-9). Results describing corticosterone levels in response to treatments are reported in Hanauer et al. (in prep.). Each bird was captured between 7 am and 12 pm, and a baseline blood sample (100 ul) was taken from the brachial vein within 4 min of capture using a 26-gauge needle and microcapillary tubes. After the sample was taken, each bird was kept in a brown paper bag for 30 min, a standard acute handling stressor that is widely used to assess physiological stress response (Sheriff et al., 2011). After 30 min in the paper bag, each bird was bled again (100 ul) to measure acute stressor-induced hormone levels. Blood was stored at 4°C until centrifugation. Later the same day, we centrifuged the microcapillary tubes to separate plasma and measure hematocrit, following which plasma was removed using Hamilton syringe and stored in -20°C until analysis.

### 2.4. GnRH injection following an acute stressor

To assess the ability of gonads to increase T, during the last sampling round at the end of the experiment (February 23-24) we injected birds with 50 ml of 25 ng/ul gonadotropin releasing hormone (GnRH, in 1M PBS, American Peptide Company Inc., Sunnyvale, CA, product no. 54- 8-23) 35 min after capture (i.e. after the stress-induced 30 min blood sample was collected). GnRH is the top regulator of testosterone synthesis that is naturally produced in hypothalamus (Ciechanowska et al., 2010). GnRH-induced testosterone levels have been shown to be repeatable (Jawor et al., 2006) and are linked to a higher probability of survival (McGlothlin et al., 2010) and morphological characteristics (Atwell et al., 2014). To administer GnRH, we cleaned the pectoral muscle using an alcohol swab, and injected 50 ul of GnRH using a Hamilton syringe. GnRH solutions were kept on ice before the injection to prevent degradation of the peptide. We then collected blood (100 ul) 30 min after injection to measure the capacity of gonads to secrete testosterone following an acute stressor.

### 2.5. Morphological measurements

After blood collection we used a ruler to measure the size of the cloacal protuberance by measuring its length from body to the cloacal opening to the nearest 0.5 mm. We also measured the pectoral muscle condition, fat score, and mass. The effects of the treatment on condition, fat, and mass are reported elsewhere (Hanauer et al. in prep).

### 2.6. Hormone assays

We measured testosterone levels in the blood plasma using commercial enzyme immune assay (EIA) from Enzo Life Sciences (Farmingdale, NY, product no. ADI-901-65) that has been previously validated for use in this species (Clotfelter et al., 2004). Testosterone was extracted from 20 ul of plasma using diethyl ether. Tritiated testosterone was added to the sample to estimate extraction efficiency (average 94.8%). Extracted hormone was reconstituted in 50 ul of 98% ethanol, followed by 300 ul of assay buffer. We followed manufacturer’s instructions for the remaining procedures. We estimated hormone concentrations in reference to a 9-point standard curve using a cure-fitting program (Microplate Manger, Bio-Rad Laboratories, Hercules, CA). Samples were distributed randomly across and within plates, and all samples and standard curves were run in duplicate. Within-plate variation was 7.01% and between-plate variation was 1.89%. We did not correct for either extraction efficiency or across-plate variation.

### 2.7 Tissue collection

One to two days after the final blood sampling (Feb 25-26), birds were euthanized using isoflurane overdose in a two-way factorial design: birds from both chronic disturbance and control treatments were euthanized either immediately after capture or after 90 min of being held in a paper bag (acute handling stressor). Literature on immediate early gene (IEG) expression and half-life suggests that 90 min following stimulus allows to capture both induction as well as reduction in IEG expression (Maney and Goodson, 2011). Since IEGs include transcription factors, we estimated that gene expression 90 min after capture was likely to capture both up-regulation and down-regulation of components of the gonadal transcriptome compared to expression immediately following capture. Gonads were dissected and flash-frozen on pulverized dry ice. To assess gonadal size, we measured gonadal mass (to the nearest mg) before RNA extraction.

### 2.8. Microarray hybridization and analyses

RNA was extracted from gonads using Trizol method (Invitrogen, Carlsbad, CA) and quantified using the Nanodrop ND-2000 (ThermoScientific, Waltham, MA). Sample integrity was verified using Agilent Bioanalyzer or TapeStation (Agilent Technologies, Santa Clara, CA). Total RNA was converted to double stranded cDNA using the TransPlex Complete Whole Transcriptome Amplification Kit (Sigma-Aldrich, St. Louis, MO, product no. WTA2) according to manufacturers instructions. We used a custom Nimblegen 12-plex microarray (Roche Nimblegen, Madison, WI) for the Dark-eyed junco (Peterson et al., 2014) to analyze the effect of treatments on transcriptome (see Supplementary Material for more detailed molecular methods). The microarray contained 33,545 assembled sequencing reads (contigs) in triplicate covering 22,765 putative genes (isogroups), which were based on Dark-eyed Junco transcriptome sequencing (Peterson et al., 2012). cDNA was hybridized to the microarray using a full round-robin design.

A series of pair-wise comparisons were tested to identify significant differences in gene expression using R package limma (Smyth, 2005) (i.e. we compared chronic disturbance vs. control within each acute handling treatment and pooled; and acute handing vs. unhandled control within each chronic treatment and pooled). We calculated a global false-discovery rate across all comparisons and used a q-value threshold of 0.05 for significance (see Peterson et al. (2014) for further details). We used topGO (Alexa and Rahnenfuhrer, 2010) with the weight algorithm (Alexa et al., 2006) to identify the gene ontology (GO) terms (Ashburner et al., 2000) that were significantly over-represented among the significantly differentially expressed genes in each comparison.

### 2.9. qPCR

The microarray did not include all of our candidate genes of interest. Therefore, we subsequently performed quantitative polymerase chain reaction (qPCR) to test our a priori predictions about genes related to testosterone synthesis and gonadal function. Specifically, we quantified expression of steroid synthesis genes (luteinizing hormone receptor (LHR), steroidogenic acute regulatory protein (StAR), cytochrome P450 side-chain cleavage (P450), cytochrome P450 17α-hydroxylase (CYP17), 3β-hydroxysteroid dehydrogenase/isomerase (3βHSD), and 17β-hydroxysteroid dehydrogenase (17βHSD)), as well as genes that encode receptors for hormones that may regulate gonadal function (sperm production: follicle-stimulating hormone receptor (FSHR); response to stressors: glucocorticoid receptor (GR), mineralocorticoid receptor (MR), gonadotropin inhibitory hormone receptor (GnIHR)). Most of the primers, except 17βHSD and FSHR primers, had been previously validated in our system (Bergeon Burns et al., 2014; Rosvall et al., 2016a; Rosvall et al., 2016b). 17βHSD and FSHR primers were designed using Zebra finch and White-throated sparrow genomes, respectively, using Primer-BLAST (Ye et al., 2012). Primer sequences and efficiencies are reported in the Supplementary Material (**Table S1**). We ran qPCR reactions using PerfeCTa SYBR Green SuperMix (Quantabio, Beverly, MA, product no. 95054) on the Roche LightCycler 480 platform (Roche Holding AG, Basel, Switzerland) using the same cDNA samples from the microarray. The cDNA concentration was similar between samples (17 to 22 ng/ul). We analyzed samples in triplicate, calculated the relative gene expression in each sample in reference to a pooled standard, and normalized this value against average expression of two housekeeping genes (Peptidylprolyl Isomerase A (PPIA) and Ribosomal Protein L4 (RPL4)) using LightCycler 480 software (Roche Holding AG, release no. 1.5.1.62). PPIA and RPL4 are two of the most stable housekeeping genes in testes of passerines (Zinzow-Kramer et al., 2014), and their expression did not differ between chronic or acute treatments. Primers for PPIA and RPL4 were designed using White-throated sparrow genome. Additional details for qPCR can be found in the Supplementary Material.

### 2.10. Statistical analysis

We analyzed our data in R (R Core Team, 2013). When running parametric tests, we confirmed that residuals satisfied the expectation of normality. If model residuals were not normal, we transformed the data. We used linear (LM) and linear mixed effects models (LMM, package nlme, Pinheiro et al. 2015) with marginal sums of squares to analyze the effect of chronic and acute treatments on reproductive physiology (hormone levels) and morphology (testes mass and CP size). We included time of sampling as a covariate in all models investigating hormone levels. When testing for change in hormone levels between sampling rounds (beginning vs end of the experiment) or the effect of acute (baseline vs acute stressor-induced), chronic (disturbance vs control) treatment, and their interaction on hormone levels, we included individual ID as random factor to account for repeated measures. If the interactions were not significant, we report the main effects from models without the interaction term. Our sample size differed between sampling rounds and between bleeds because of low blood volume in some of the samples.

We used two different multivariate approaches to analyze the effect of treatments on candidate gene expression (determined by qPCR). Since the expression of steroidogenic genes showed significant pairwise correlations, we used principal components analysis to summarize the covariation in gene expression by creating independent principal components. We then used the first principal component as a dependent variable in ANOVA to ask if chronic or acute treatments, or their interaction affected steroidogenic gene expression. Following this, we used ANOVAs to analyze each gene separately and corrected for multiple comparisons using the Benjamini-Hochberg (BH) method.

Because the expression of receptors that may downregulate gonadal function (GR, MR, and GnIHR) was not correlated to each other and they mediate different functions in animals, we used MANOVA to analyze the effect of chronic and acute treatments on the overall sensitivity to stress-signaling. Following MANOVA, we analyzed each gene independently using ANOVA and corrected for multiple comparisons using the Benjamini-Hochberg (BH) method. FSHR expression was analyzed independently using a linear model, because its function is not know to relate to stress-signaling or steroidogenesis.

## 3. RESULTS

### 3.1. Testosterone

#### Pre-treatment

At the outset of the experiment neither baseline testosterone (LM, n=20, F_2,17_=0.086, p=0.773) nor testosterone after acute handling stressor (LM, n=14, F_2,11_=1.207, p=0.295) differed between the chronic and control treatment groups, and acute handling stressor did not reduce testosterone compared to baseline levels (LMM, n=33, F_1,10_=0.226, p=0.645, **Figure 2A**). Baseline testosterone increased throughout the experiment in both treatments (LMM, n=46,F_1,12_=15.716, p=0.002), consistent with recrudescence of gonads in response to increasing day length.

**Figure 2.**
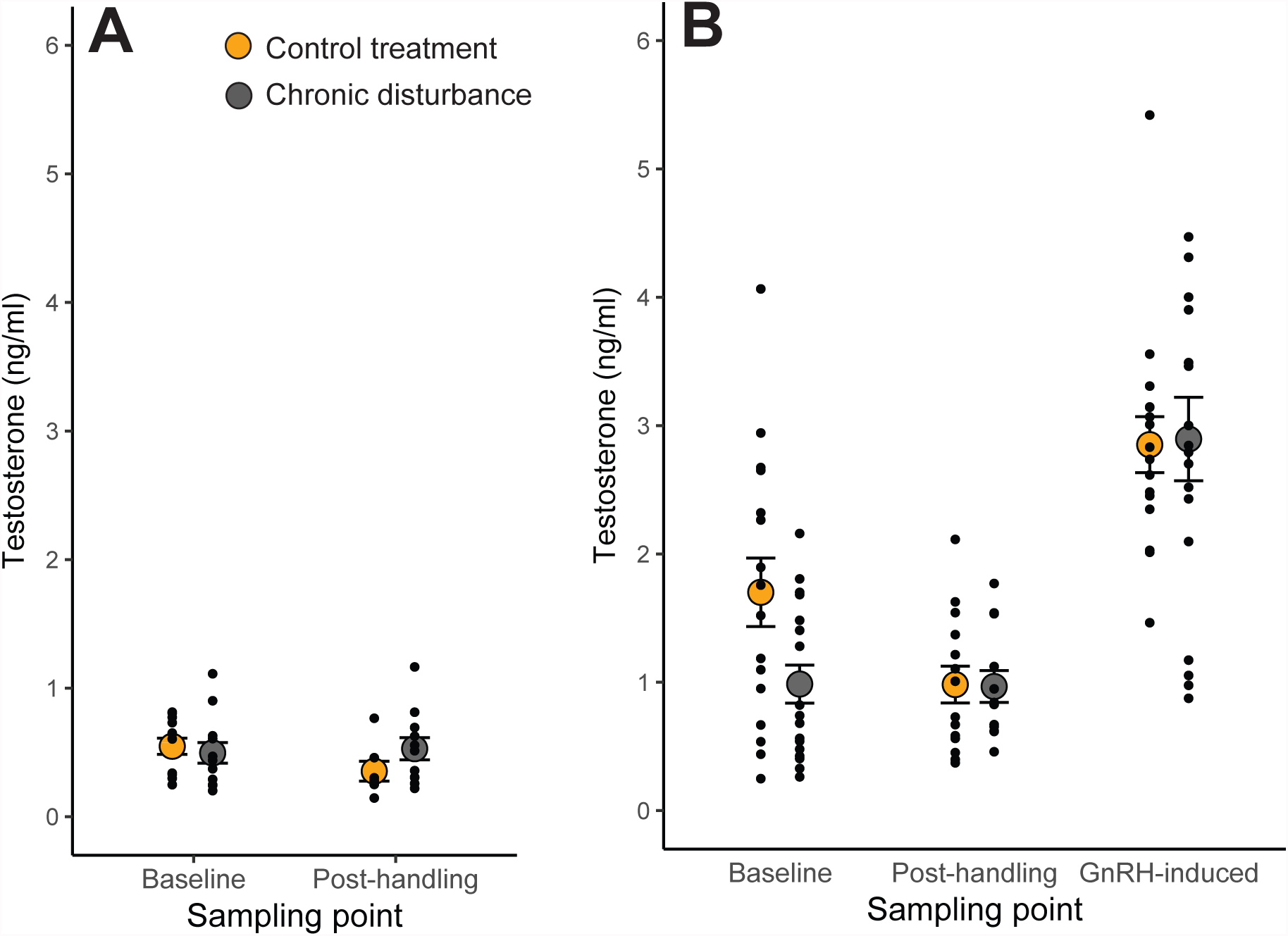
Effect of chronic disturbance and acute stressor on testosterone levels. A) testosterone levels before the experiment; B) testosterone levels after after the experiment. Shown are baseline testosterone levels, post-handling levels, and GnR-Hinduced testosterone levels. GnRH-induced testosterone was not measured before the experiment.

#### Post-treatment

After three weeks of treatment, birds from the chronic disturbance group had marginally lower baseline testosterone levels than controls (LM, n=27,F_2,24_=3.643, p=0.068). Testosterone levels after 30 min of acute handling stressor did not differ between treatments (LM, n=21, F_2,18_= 0.584, p=0.455). There was a significant interaction between the two long-term treatments (chronic and control) and acute treatments (handled vs unhandled) on testosterone levels (LMM, n=48, F_1,17_=5.935, p=0.026, **Figure 2B**). Post-hoc ANOVAS showed that birds from the control treatment showed a significant reduction in their testosterone levels from baseline to post-acute handling stress (LMM, n=24, F_1,8_=5.842, p=0.042), while birds from the chronic disturbance treatment did not (LMM, n=24, F_1,9_=0.793, p=0.396). Control and chronically disturbed birds significantly increased their testosterone in response to GnR-Hinjection compared to pre-injection levels (LMM, n=50,F_1,19_=63.527, p<0.001), but the GnR-Hinduced testosterone did not differ between treatment groups (LMM, n=50, F_1,18_=0.274, p=0.607, **Figure 2C**).

### 3.2. Reproductive organs and relationship with physiology

Testes mass was not affected by the chronic disturbance treatment (LM, n=35, F_1,33_=0.873, p=0.357, **Figure 3A**). Testes mass was positively related to testosterone levels after acute handling stress (LM, n=21, F_2,18_=6.620, p=0.019) as well as GnRH-induced testosterone levels (LM, n=29, F_2,26_=8.292, p=0.008), but was not related to baseline testosterone (LM, n=27, F_2,24_=0.435, p=0.516). The relationship between testes mass and testosterone was not dependent on the chronic treatment (interaction between gonad mass and treatment on post-handling T: LM, n=21, F_4,16=0.025_, p=0.877; on GnRH-induced T: LM, n=29, F_4,24_=1.201, p=0.284). Cloacal protuberance (CP) size showed significant differences between treatments, with chronically disturbed birds having larger CPs than control birds (LM, n=33, F_1,31_=22.198, p<0.001, **Figure 3B**).

**Figure 3:**
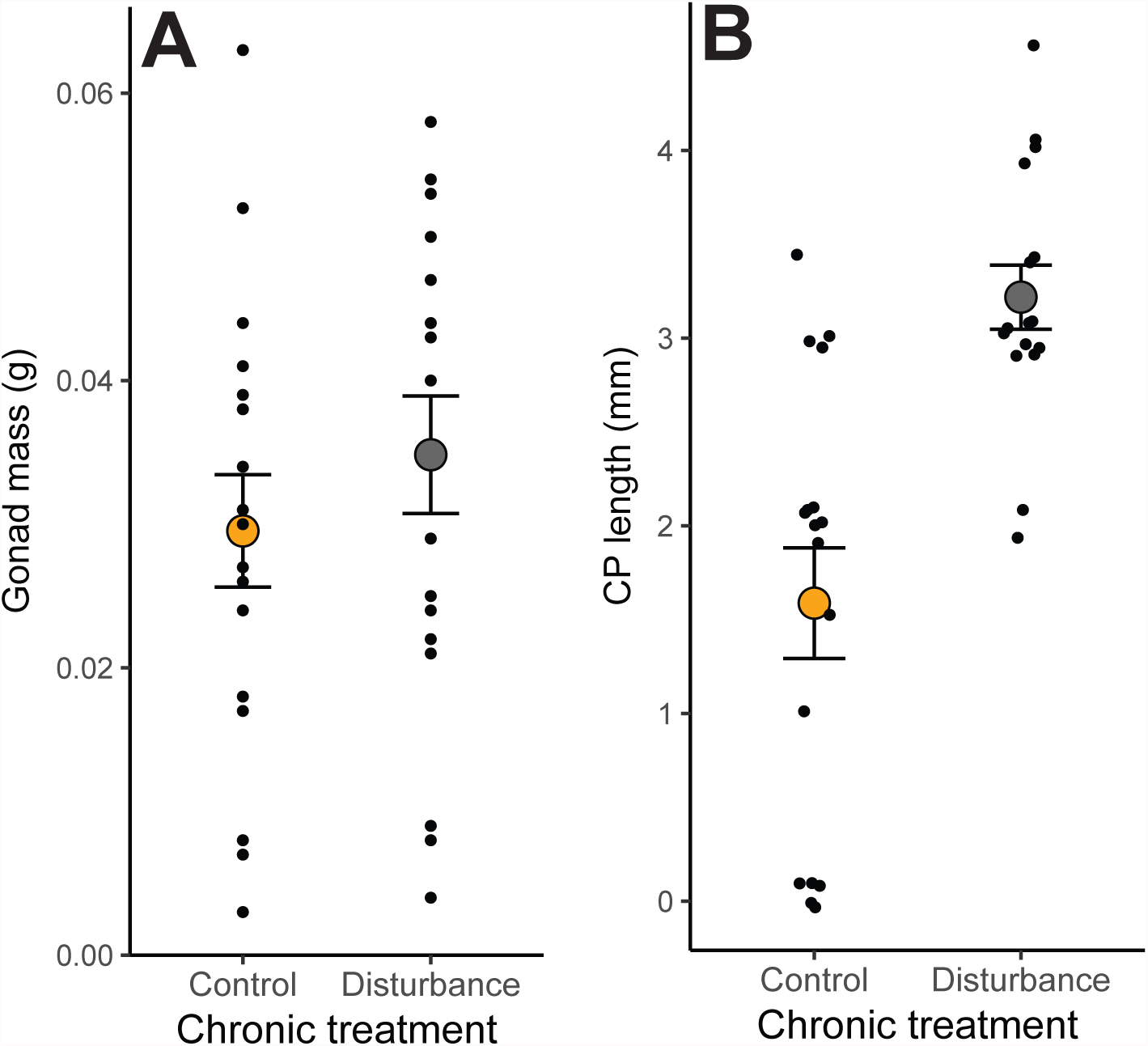
Effect of chronic disturbance on reproductive organs. A) Gonad mass; B) Cloacal protuberance (CP). Points are jittered in B to show sample size.

### 3.3. Gonadal transcriptome

We found that the expression of 16 transcripts were significantly affected by the chronic disturbance treatment after correcting for false discovery rate (out of 20390 total expressed in gonads in this study, 0.078%; **Table S2**). GO analysis identified 3 terms that were overrepresented among the significantly differentially expressed genes. These included: electron transport chain, extracellular structure organization (biological process), and glycosaminoglycan binding (molecular function) (**Table 1**). None of the genes that were differentially expressed between tissues in response to chronic treatment are known to link clearly to the stress response or reproductive function.

**Table 1:**
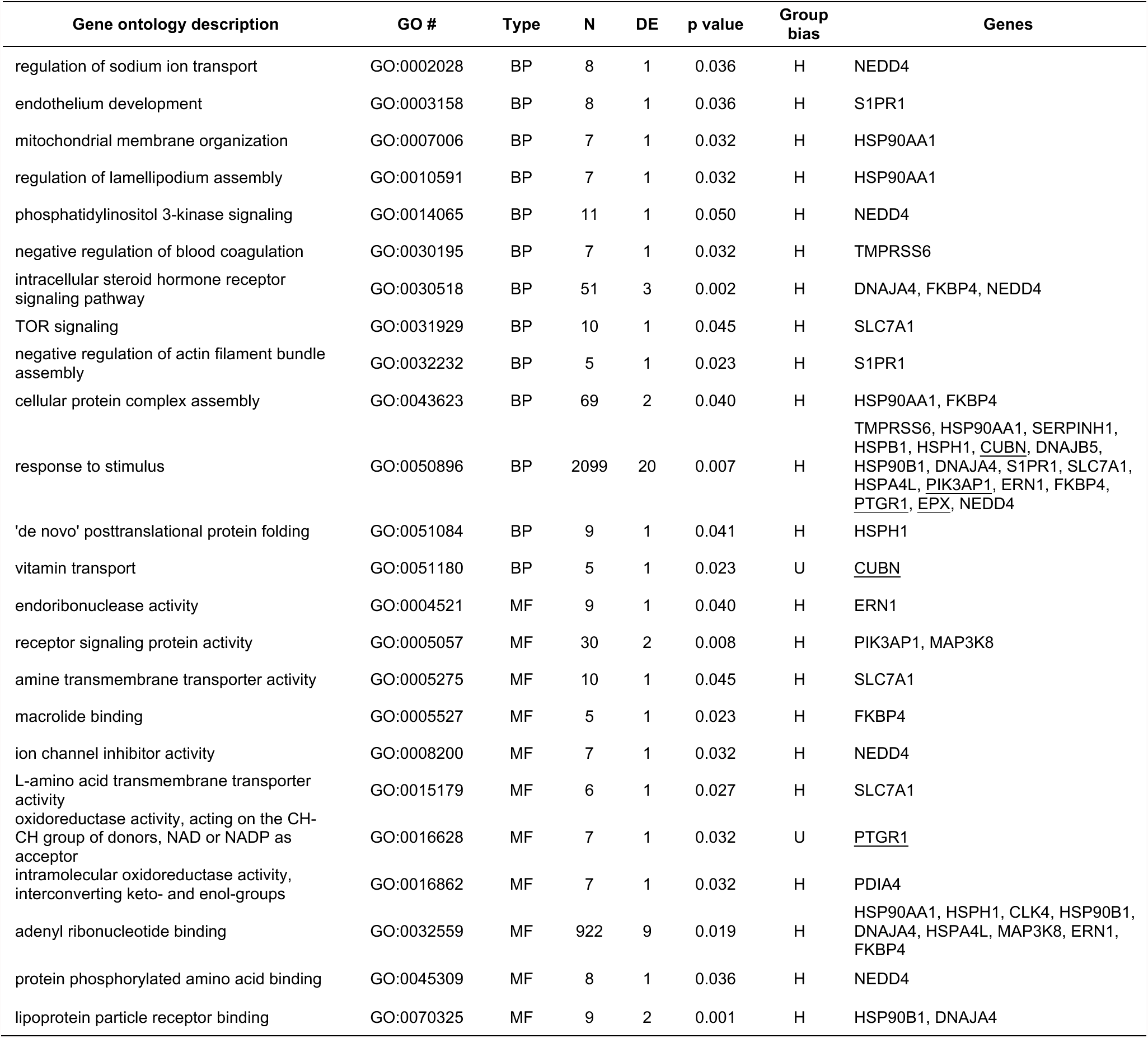
Gene ontology (GO) processes significantly affected by the acute handling treatment (H) compared to unhanded control treatment (U). Type= GO category; BP = biological process, MF = molecular function; N = number of genes per GO category in microarray; DE = number of genes per GO category significantly differentially expressed between treatments. Non-underlined genes were up-regulated in the H treatment. Underlined genes were up-regulated in C treatment. Group bias refers to direction of upregulation.

| Gene ontology description | GO # | Type | N | DE | p value | Group bias | Genes |
| --- | --- | --- | --- | --- | --- | --- | --- |
| regulation of sodium ion transport | GO:0002028 | BP | 8 | 1 | 0.036 | H | NEDD4 |
| endothelium development | GO:0003158 | BP | 8 | 1 | 0.036 | H | S1PR1 |
| mitochondrial membrane organization | GO:0007006 | BP | 7 | 1 | 0.032 | H | HSP90AA1 |
| regulation of lamellipodium assembly | GO:0010591 | BP | 7 | 1 | 0.032 | H | HSP90AA1 |
| phosphatidylinositol 3-kinase signaling | GO:0014065 | BP | 11 | 1 | 0.050 | H | NEDD4 |
| negative regulation of blood coagulation | GO:0030195 | BP | 7 | 1 | 0.032 | H | TMPRSS6 |
| intracellular steroid hormone receptor signaling pathway | GO:0030518 | BP | 51 | 3 | 0.002 | H | DNAJA4, FKBP4, NEDD4 |
| TOR signaling | GO:0031929 | BP | 10 | 1 | 0.045 | H | SLC7A1 |
| negative regulation of actin filament bundle assembly | GO:0032232 | BP | 5 | 1 | 0.023 | H | S1PR1 |
| cellular protein complex assembly | GO:0043623 | BP | 69 | 2 | 0.040 | H | HSP90AA1, FKBP4 |
| response to stimulus | GO:0050896 | BP | 2099 | 20 | 0.007 | H | TMPRSS6, HSP90AA1, SERPINH1, HSPB1, HSPH1, <u>CUBN</u>, DNAJB5, HSP90B1, DNAJA4, S1PR1, SLC7A1, HSPA4L, PIK3AP1, ERN1, FKBP4, <u>PTGR1</u>, <u>EPX</u>, NEDD4 |
| 'de novo' posttranslational protein folding | GO:0051084 | BP | 9 | 1 | 0.041 | H | HSPH1 |
| vitamin transport | GO:0051180 | BP | 5 | 1 | 0.023 | U | <u>CUBN</u> |
| endoribonuclease activity | GO:0004521 | MF | 9 | 1 | 0.040 | H | ERN1 |
| receptor signaling protein activity | GO:0005057 | MF | 30 | 2 | 0.008 | H | PIK3AP1, MAP3K8 |
| amine transmembrane transporter activity | GO:0005275 | MF | 10 | 1 | 0.045 | H | SLC7A1 |
| macrolide binding | GO:0005527 | MF | 5 | 1 | 0.023 | H | FKBP4 |
| ion channel inhibitor activity | GO:0008200 | MF | 7 | 1 | 0.032 | H | NEDD4 |
| L-amino acid transmembrane transporter activity | GO:0015179 | MF | 6 | 1 | 0.027 | H | SLC7A1 |
| oxidoreductase activity, acting on the CH-CH group of donors, NAD or NADP as acceptor | GO:0016628 | MF | 7 | 1 | 0.032 | U | <u>PTGR1</u> |
| intramolecular oxidoreductase activity, interconverting keto- and enol-groups | GO:0016862 | MF | 7 | 1 | 0.032 | H | PDIA4 |
| adenyl ribonucleotide binding | GO:0032559 | MF | 922 | 9 | 0.019 | H | HSP90AA1, HSPH1, CLK4, HSP90B1, DNAJA4, HSPA4L, MAP3K8, ERN1, FKBP4 |
| protein phosphorylated amino acid binding | GO:0045309 | MF | 8 | 1 | 0.036 | H | NEDD4 |
| lipoprotein particle receptor binding | GO:0070325 | MF | 9 | 2 | 0.001 | H | HSP90B1, DNAJA4 |

Acute handling stressor caused significant change in the expression of 168 transcripts compared to unhandled controls (out of 20390 total expressed in gonads, 0.823%; **Table S3**). Among genes that were contributing to these terms were a variety of heat shock proteins (HSPB1, DNAJA4, HSPA4L, HSP90AA1), and genes associated with inflammation and cytokine signaling (IL4R, PIK3AP1, MAP3K8). GO analysis identified a number of terms that were overrepresented in the set of genes that showed significant treatment effect including: receptor signaling protein activity, intracellular steroid hormone receptor signaling pathway, cellular protein complex assembly, response to stimulus, and ‘de novo’ posttranslational protein folding (**Table 2**). One unannotated transcript with an unknown function showed a significant interaction between the long-term treatments (chronic vs control) and acute treatments (handled vs unhandled).

**Table 2:** Gene ontology processes significantly affected by chronic disturbance vs control treatments. N = number of genes per GO category in microarray; DE = number of genes per GO category significantly differentially expressed between treatments. C = control treatment.

| Gene ontology description | GO # | Type | N | DE | p value | Group bias | Genes |
| --- | --- | --- | --- | --- | --- | --- | --- |
| electron transport chain | GO:0022900 | BP | 37 | 1 | 0.023 | C | NDUFA3 |
| extracellular structure organization | GO:0043062 | BP | 81 | 1 | 0.049 | C | POSTN |
| glycosaminoglycan binding | GO:0005539 | MF | 48 | 1 | 0.027 | C | POSTN |

### 3.4. Testosterone synthesis and HPG axis receptors

Principal components analysis of six genes involved in testosterone synthesis (LH-R, STAR, p450scc, CYP-17, 3bHSD, 17bHSD) showed that PC1 loaded negatively for all genes and explained 45% of the variation in expression (**Table S4**). This suggests that the expression of the testosterone synthesis genes co-varied in the same direction between individuals. However, variation in PC1 was not explained by long-term disturbance (LM, n=35, F_3,31_=0.261, p=0.613), acute handling stressor (LM, n=35,F_3,31=_ 1.700, p=0.202), or the interaction of these treatments (LM, n=35,F_3,31_=0.231, p=0.634). Since testosterone levels decreased significantly in response to acute handling stressor only in the control treatment, we conducted a separate PCA of only the control individuals (see **Table S5** for loadings). This analysis showed that acute handling stressor tended to reduce the expression of testosterone synthesis genes 90 min later, although this effect was not significant (LM, n=17,F_1,15_=3.157, p=0.096). Analysis of each of the genes individually using ANOVA, showed that CYP17 levels were marginally lower in handled control birds (LM, n=17,F_1,15_=4.418, p=0.053, see **Table S6** for other genes) although this effect was not significant after multiple comparison correction (p=0.317, **Figure S1**).

Neither handling (LM, n=35,F_2,32_= 2.051, p=0.162), nor chronic disturbance (LM, n=35, F_2,32_ = 2.733, p= 0.108), nor their interaction (LM, n=35,F_3,31=_ 0.437, p= 0.514) affected expression of FSH-R (**Figure S2**).

### 3.5. Gonadal sensitivity to regulation by HPA

MANOVA showed that acute handling stressor (n=35, Wilks=0.676,F_3,30_=4.791, p=0.008), but not chronic disturbance (n=35, Wilks=0.907, F_3,30_=1.027, p=0.395), nor interaction between chronic treatment and acute handling stressor (n=35, Wilks= 0.855, F_3,29_=1.643, p=0.201) affected expression of receptors for hormones that are known to suppress or inhibit testosterone production. Post-hoc linear models comparing handled to unhandled birds showed that acute handling stressor significantly down-regulated GR mRNA expression (LM, n=35, F_2,32_=5.207, p=0.029, **Figure 4A**) and marginally up-regulated of GnIHR expression (LM, n=35, F_2,32_=3.517, p=0.070, **Figure 4C**), although these effects were not significant after correction for multiple comparisons (see **Table S7**).

**Figure 4.**
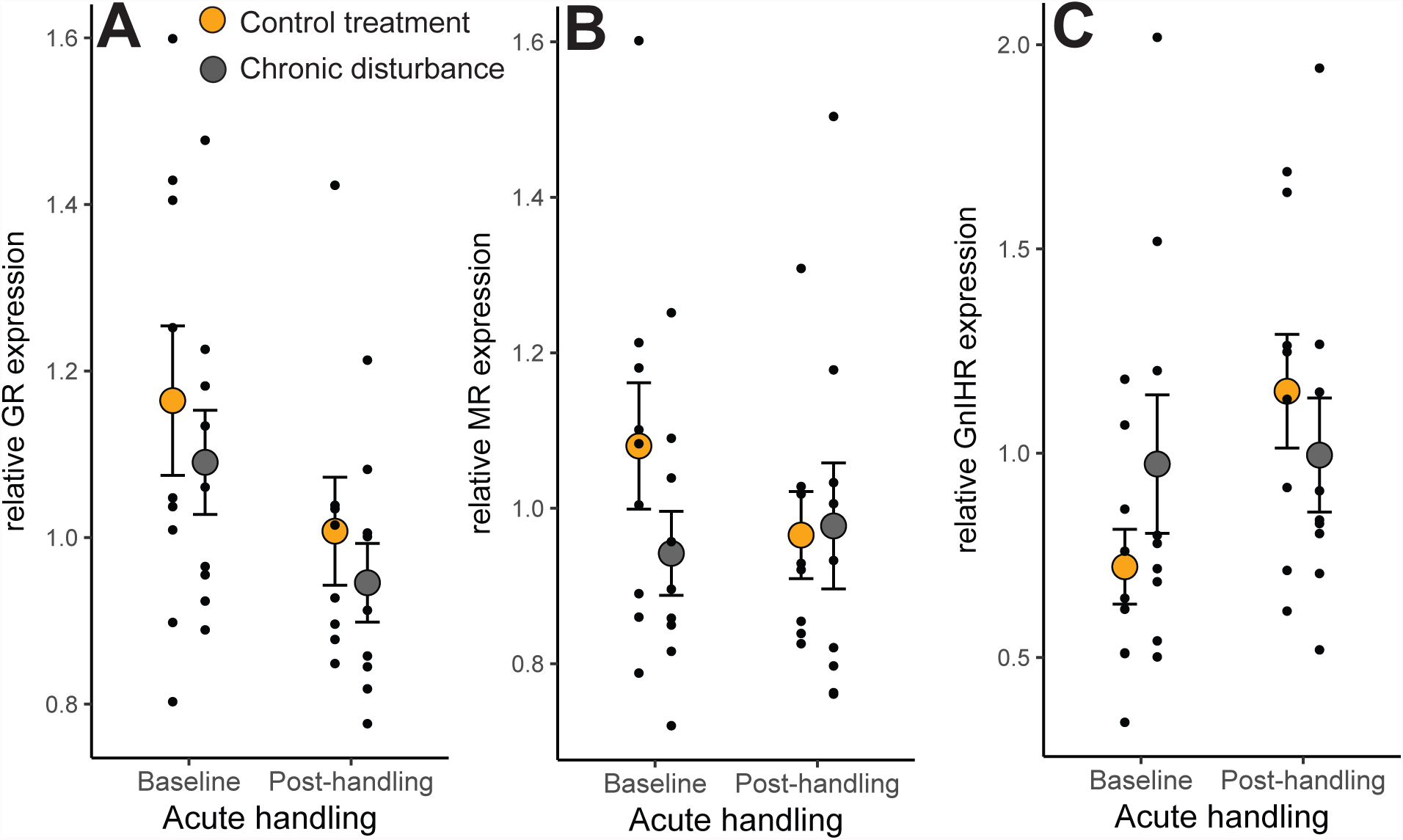
Effect of chronic disturbance and acute handling stressor on A) glucocorticoid receptor (GR) expression; B) mineralocorticoid receptor (MR) expression; C) gonadotropin inhibitory hormone receptor (GnIHR) expression in gonads.

## 4. DISCUSSION

In this study we experimentally tested how chronic disturbance, acute handling stressor, and their interaction affected gonadal function in a songbird during seasonal gonadal recrudescence. We showed that birds in the chronic disturbance treatment had marginally lower baseline testosterone compared to control animals. Acute handling stressor reduced testosterone levels in control animals, but not in animals from the chronic disturbance treatment. The chronic disturbance treatment did not affect the ability to produce testosterone in response to exogenous stimulation from GnRH, and gonad size did not differ between chronic disturbance and control treatments. Surprisingly, birds in the chronic disturbance treatment had significantly larger cloacal protuberances than birds in the control treatment. Chronic disturbance had very little effect on gonadal gene expression: neither steroidogenic enzyme expression, nor expression of receptors associated with potential regulation of gonadal function showed significant differences between the chronic disturbance and control treatments. Overall, chronic disturbance had a significant effect on only a handful of genes in the transcriptome. Acute handling stressor, on the other hand, had a comparatively strong effect on the gonadal transcriptome, a marginal suppressive effect on steroidogenesis enzyme gene expression, a suppressive effect on the expression of glucocorticoid receptor mRNA, and a marginal positive effect on GnIH-R gene expression compared to unhandled animals. There was little evidence of an interaction between chronic disturbance and acute stressor on the expression of gonadal transcriptome. Collectively, these findings shed light on mechanisms by which short and long-term stressors, and their interaction, affect reproductive function.

### 4.1. Chronic stressor and gonadal function

Birds from the chronic disturbance treatment had marginally lower baseline testosterone levels compared to the control treatment. Our study adds to other reports in a variety of organisms that have shown a decrease in testosterone in response to chronic disturbance (Pickering et al. 1987; Moore et al. 1991; Retana-Márquez et al. 2003) although some studies do not show this effect (Armario and Castellanos 1984; Jones and Bell 2004). Long-term suppression of testosterone levels in response to chronic stressor has important ecological implications. Human-induced rapid environmental change is exposing animals to novel environments and rapid changes in the disturbance regimes, which have been shown to have strong negative effects on foraging and survivorship (Kerley et al., 2002; Sih et al., 2011). If a decrease in testosterone in response to chronic disturbance results in reduced investment in reproduction in general, this may exacerbate the negative effects of disturbance on fitness.

Mechanistically, the differences in testosterone levels between the chronic disturbance and control birds in this study were not explained by differences in gonadal physiology. Testes mass and GnRH-induced testosterone levels did not differ between treatments, indicating that the ability of the gonads to produce and elevate testosterone was not different. This suggests that the marginally lower testosterone levels in the chronically disturbed animals and their insensitivity to acute handling stress compared to control animals were not caused by long-lasting differences in the ability of the testes to produce testosterone.

Differences in testosterone levels instead may be mediated by local or systemic signaling that transiently suppresses or activates testosterone synthesis in the gonad or testosterone metabolism in the liver. Experimental studies have shown that GnIH (McGuire and Bentley, 2010) and corticosterone (McGuire et al., 2013) downregulate testosterone by acting directly on gonadal tissue. This effect may be mediated by either changes in the levels of these hormones or by changes in the expression of their receptors. Baseline corticosterone did not differ between chronic disturbance and control treatments (Hanauer et al. in prep.). We could not assess GnIH expression in gonads or brain, therefore we do not know if chronic stressors upregulated GnIH synthesis. Our gene expression results showed that chronic disturbance did not affect the expression of receptors for GnIH (GnIHR), corticosterone (GR, MR), or luteinizing hormone (LHR), a top regulator of testosterone synthesis. Furthermore, chronic disturbance did not have a significant effect on steroidogenic gene expression or the testicular transcriptome in general, showing that gonadal function was nearly unaffected by the chronic disturbance treatment. This suggests that local signaling at the level of the gonad is unlikely to be the cause of lower testosterone levels or the decreased sensitivity to acute handling stressor that we observed in the chronically disturbed animals. Future analysis of the post-translational effects of stressors, as well as other tissues (e.g. hypothalamus, liver) collected from these animals may elucidate the mechanisms that caused the differences in testosterone levels between treatments (Lynn et al., 2015).

A surprising finding in this study was that the cloacal protuberances (CPs) were significantly larger in the chronically disturbed birds compared to control animals. CPs in birds are sperm storage organs that develop in males during the breeding season (Ray Salt, 1954) and are regulated by testosterone (Witschi, 1961). Males with larger CPs store more sperm (Tuttle et al., 1996) and larger CPs are hypothesized to allow faster copulation (Birkhead et al., 1993). Barring spurious results, our finding may indicate that birds in the chronic disturbance group may be investing more in sperm production compared to the control animals, perhaps to advance their reproductive readiness or enable more rapid copulation in an uncertain environment. It is important to note that neither CPs nor testes reached their full size during this study (i.e testes mass was 21% of mature reproductive size (Bergeon Burns et al., 2014)), therefore our CP findings, coupled with measurement error, may reflect functionally irrelevant differences.

### 4.2. Acute stressor and gonadal function

Birds in the control treatment showed a significant decrease in testosterone levels in response to the acute handling stressor. Suppression of testosterone in response to acute stressors has been shown in many other studies (Deviche et al., 2010; Deviche et al., 2012; Moore et al., 1991). This effect may be due to reduction in testosterone synthesis, or an increase in testosterone metabolism by liver (Lynn et al., 2015). We found a marginally significant decrease in the expression of steroidogenic enzyme genes in birds that had experienced an acute handling stressor compared to birds that were unhandled. Because the expression of steroidogenic enzymes is positively correlated with testosterone levels (Rosvall et al., 2016b), decrease in expression of testosterone synthesis genes may be responsible for suppression of testosterone in response to handling.

Downregulation of steroidogenic enzyme gene expression in response to the acute handling stressor may be due to either a change in the local or systemic signaling or because of a change in sensitivity to this signaling (Ernst et al., 2015). We found a significant overall change in hormone receptor gene expression, primarily downregulation of GR expression and upregulation of GnIHR expression, in response to the acute handling stressor. This suggests that gonads in acutely stressed birds may be less sensitive to regulation by glucocorticoids, such as corticosterone, but more sensitive to another important hormone – GnIH – that has an inhibitory effect on reproductive physiology (Tsutsui et al., 2010). MR expression was not affected by acute stressor. While GR and MR are both involved in regulating the stress response (Dickens et al., 2009) they fulfill different functional roles, with the high affinity MR (Reul and Kloet, 1985) thought to be responsible regulating baseline functions when CORT levels are low, while the low-affinity GR regulates stress-response when CORT levels are high (Ulrich-Lai and Herman, 2009; Wingfield, 2012). Thus, the GR-specific reduction we observed in response to an acute handling stressor is consistent with known differences between these two types of CORT-binding receptors. We could not assess the expression of GnIH in the gonads or brain, therefore we do not know if acute or chronic stressors, in addition to GnIHR, also upregulated GnIH synthesis. In another study, we validated that corticosterone levels increased following this acute handling stressor (Hanauer et al. in prep).

Acute handling stress caused significant changes across the testicular transcriptome compared to unhandled animals. The gene ontology (GO) analysis did not indicate that the acute handling treatment affected testes-specific functions, such as spermatogenesis or steroidogenesis, although we note that several candidate genes were not present on this array. Instead, GO analysis suggested that acute handling stress induced cellular stress response, resulting in upregulation of a handful of heat shock proteins. Heat shock proteins are molecular chaperones that under normal conditions facilitate protein assembly and are upregulated in response to stressors (Akerfelt et al., 2010). Heat shock protein expression in gonads increases during spermatogenesis and oogenesis (Neuer et al., 2000) which may ensure that the development of gametes is shielded from environmental stressors. Importantly, some of the same genes (e.g. HSP90AA1, DNAJA4) upregulated in response to acute handling stress in this study, were also upregulated in chicken testes in response to heat stress (Wang et al., 2015), indicating that gonads may have a generalized cellular stress response that is upregulated in response to a variety of stressors. Analysis of transcriptomic changes in other tissues will indicate if the change in gonadal transcriptome represents testes-specific response to stress, or if response to stress between tissues differs.

### 4.3. Interaction between chronic and acute stressors

Birds from chronic disturbance and control treatments responded differently to the acute handling stressor: whereas handling reduced testosterone levels in control birds, it did not affect testosterone levels in birds from the chronic disturbance treatment. This difference can be interpreted in two non-exclusive ways.

First, the reduced impact of the acute stressor on testosterone in birds from the chronic disturbance group might have arisen because their reproductive function was already downregulated to a degree that prohibited further decrease (“floor effect” *sensu* (Sapolsky et al., 1984)). In our study, however, both the baseline and post-handling testosterone levels of chronically disturbed animals were significantly higher than testosterone levels at the beginning of the experiment. This suggests that, mechanistically, testosterone levels could have shown a further decrease below the levels observed in response to the acute handling stressor. It is possible, however, that the lack of decrease in testosterone in response to handling in chronically disturbed birds may be due to reduced testosterone metabolism by the liver. If this was the case, we would predict that acute stressors would start suppressing testosterone levels later in chronically disturbed animals compared to control animals. Unfortunately, our experimental design did not allow us to test this possibility due to the GnRH injection during blood sampling.

Second, the difference in the effect of handling on testosterone levels between chronic disturbance and control treatments could be explained by a lower sensitivity of the chronically disturbed birds to stressors compared to the control individuals. We did not find any effect of chronic disturbance on gonadal signaling that would explain this difference, suggesting that differences in sensitivity to stress may exist in the HPG axis tissues that are upstream from the gonad (pituitary, hypothalamus), or other hormonal targets that interact with testosterone production.

From an ecological perspective, the first alternative (floor effect) could be interpreted as a consequence of homeostatic overload (Romero et al., 2009). Homeostatic overload refers to the cases where overexposure to stressors, and the associated increase in the frequency of stress response, results in a pathological state that leads to compromised organismal function (Romero et al., 2009). In our study, the high frequency of stressors may have have resulted in a compromised ability to produce testosterone. However, because birds from both chronic and control treatments elevated their testosterone in response to GnRH to the same degree, this explanation seems unlikely. Instead, the floor effect could represent be a result of a “best of a bad job” strategy, wherein individuals of low quality or in a bad environment maximize their fitness by playing it safe (Sih and Bell, 2008). In this case, animals may opt to keep the resources diverted from reproduction to maximize their ability to respond to stressors. This strategy would be more likely to be adaptive in long-lived animals and in environments were the disturbance frequency is likely to change. However, even in short-lived animals, strategies with low investment in reproduction can have higher fitness compared to cases when animals invest in reproduction at the cost of survival (e.g. sneaky copulations by alternative morphs in beetles (Moczek and Emlen, 2000)).

The second alternative (differences in sensitivity to stress), is consistent with the life history prediction that under conditions of low expected future reproductive success, stress should have a less negative effect on reproduction compared to situations where expected reproductive success is high (Wingfield and Sapolsky, 2003). Animals that develop in environments with high frequency of stressors may change their physiology to reduce sensitivity of HPG axis to stress, thus allowing them to maintain reproductive function during stressful episodes, despite the possible cost to self-maintenance (Wingfield and Sapolsky, 2003). This prediction has been supported by a study in a natural system (Abolins-Abols et al., 2016), where the reproductive phenotype of urban animals is less sensitive to stressors compared to rural animals.

In contrast to our testosterone findings, we did not find and an interaction between chronic and acute treatments on the gonadal transcriptome, although we note that the marginal effects of the chronic treatment reduce our power to explore this interaction. This finding nevertheless provides further evidence that chronic disturbance does not affect gonadal capacity to produce testosterone or respond to stress. Instead, we hypothesize that the differential effect of handling stress on testosterone levels in the chronic disturbance and control treatments may be due to differences in upstream signaling from the pituitary or hypothalamus, which regulates the activity of the molecular machinery in the gonad. Further work will be needed to explore alternatives, such as post-translational effects of stress that are not captured at the level of gene expression, or temporal dynamics wherein acute and chronic stressors interact at only specific time-points following a stressor. Despite these uncertainties, our results nevertheless demonstrate that the gonad is relatively robust to interacting effects of acute and chronic stressors on gene expression.

### 4.4. Summary

Overall our results show that chronic and acute stressors suppress testosterone release by the gonads, but that the effect of acute stressors differs depending on the frequency of stressors in the environment. These differences in gonadal function between disturbance environments are unlikely to be mediated by changes in gonadal gene expression, but are more likely to reside upstream of the gonad. Whereas chronic treatment had a negligible effect on the gonadal transcriptome, acute handling stress significantly upregulated major components of cellular stress response and affected the expression of hormone receptors involved in downregulation of testosterone production. These results suggest a potential mechanism for regulating testosterone decrease in response to acute stressors. Furthermore, they show that transient changes in gene expression in response to acute stressors are different from more permanent responses to chronic stressors. An important future direction therefore is to identify the mechanisms responsible for differences in testosterone levels between animals experiencing different disturbance regimes.

This study is among the first to experimentally test the mechanisms by which acute and chronic stressors interact to influence reproduction. We demonstrate patterns that, adaptive or not, are doubtlessly important in understanding how animals respond to chronic and acute stressors. We therefore urge further study on the adaptive significance and mechanisms that mediate the effect of chronic stressors on testosterone levels and other reproductive functions.

## Acknowledgments

This study was conducted in accordance with Indiana University Animal Care and Use Committee guidelines, protocol #12-050-08. We thank A. Kimmitt, S. Slowinski, H. Kassab, A. Brenner, A. McKay, C. Taylor, and N. Fletcher for assistance with disturbance manipulations, animal care, and sampling. We thank L. Sloan, who extracted RNA, M. Stephens, who ran the microarray at the University of Notre Dame Genomics & Bioinformatics Core Facility, and D. Rusch and A. Fellows from IU’s Center for Genomics and Bioinformatics for assistance with microarray data analysis. We thank C. Bergeon-Burns for providing us with primer sequences and advice on lab work, R.A. Stewart for assistance with molecular sex determination, and the Center for the Integrative Study of Animal Behavior at IU for providing laboratory facilities and equipment.

## Data availability

Data will be published in the Dryad Digital Repository upon acceptance.

